# Reshuffling yeast chromosomes with CRISPR/Cas9

**DOI:** 10.1101/415349

**Authors:** Aubin Fleiss, Samuel O’Donnell, Téo Fournier, Nicolas Agier, Stéphane Delmas, Joseph Schacherer, Gilles Fischer

## Abstract

Genome engineering is a powerful approach to study how chromosomal architecture impacts phenotypes. However, quantifying the fitness impact of translocations independently from the confounding effect of base substitutions has so far remained challenging. We report a novel application of the CRISPR/Cas9 technology allowing to generate with high efficiency both uniquely targeted and multiple concomitant reciprocal translocations in the yeast genome. Targeted translocations are constructed by inducing two double-strand breaks on different chromosomes and forcing the trans-chromosomal repair through homologous recombination by chimerical donor DNAs. Multiple translocations are generated from the induction of several DSBs in LTR repeated sequences and promoting repair using endogenous uncut LTR copies as template. All engineered translocations are markerless and scarless. Targeted translocations are produced at base pair resolution and can be sequentially generated one after the other. Multiple translocations result in a large diversity of karyotypes including in some instances large segmental duplications. To test the phenotypic impact of translocations, we first recapitulated in a lab strain the *SSU1/ECM34* translocation providing increased sulphite resistance to wine isolates. Surprisingly, the same translocation in a laboratory strain resulted in decreased sulphite resistance. However, adding the repeated sequences that are present in the *SSU1* promoter of the resistant wine strain induced sulphite resistance in the lab strain, yet to a lower level than that of the wine isolate, implying that additional polymorphisms also contribute to the phenotype. These findings illustrate the advantage brought by our technique to untangle the phenotypic impacts of structural variations from confounding effects of base substitutions. Secondly, we showed that strains with multiple translocations display large phenotypic diversity in a wide range of environmental conditions. No coding sequence or promoter region was altered by the multiple translocations showing that simply reconfiguring chromosome architecture is sufficient to provide fitness advantages in stressful growth conditions.

**AUTHOR SUMMARY:** Chromosomes are highly dynamic objects that often undergo large structural variations such as reciprocal translocations. Such rearrangements can have dramatic functional consequences, as they can disrupt genes, change their regulation or create novel fusion genes at their breakpoints. For instance, 90-95% of patients diagnosed with chronic myeloid leukemia carry the Philadelphia chromosome characterized by a reciprocal translocation between chromosomes 9 and 22. In addition, translocations reorganize the genetic information along chromosomes, which in turn can modify the 3D architecture of the genome and potentially affect its functioning. Quantifying the fitness impact of translocations independently from the confounding effect of base substitutions has so far remained challenging. Here, we report a novel CRISPR/Cas9-based technology allowing to generate with high efficiency and at a base-pair precision either uniquely targeted or multiple reciprocal translocations in yeast, without leaving any marker or scar in the genome. Engineering targeted reciprocal translocations allowed us for the first time to untangle the phenotypic impacts of large chromosomal rearrangements from that of point mutations. In addition, the generation of multiple translocations led to a large reorganization of the genetic information along the chromosomes. Although no gene was disrupted, we showed that solely shuffling the genome resulted in the emergence of fitness advantage in stressful environmental conditions.

## INTRODUCTION

Genetic polymorphisms are not restricted to base substitutions and indels but also include large-scale Structural Variations (SVs) of chromosomes. SVs comprise both unbalanced events, often designated as copy number variations (CNVs) including deletions and duplications, and balanced events that are copy number neutral and include inversions and translocations. Both have a phenotypic impact, however the prevalence and the fitness effect of balanced SVs has been less documented than CNVs, partly because they are much more challenging to map than CNVs and also because quantifying their fitness contribution independently from the confounding effect of base substitutions remains challenging. Natural balanced chromosomal rearrangements result from the exchange of DNA ends during the repair of Double Strand Breaks (DSBs) either through Homologous Directed Repair (HDR) between dispersed repeats or intact chromosomes carrying internal repeat sequences homologous to the DNA ends [1,2] or through Non-Homologous End Joining (NHEJ) [3]. Artificial balanced rearrangements are classically engineered by inducing targeted DSBs and promoting repair through both HDR and NHEJ. However, inducing targeted DSBs and engineering scar-less chromosomal rearrangements has remained difficult. In early studies structural variants were obtained through I-SceI-induced DSB repair between split alleles of a selection marker [4,5]. In later developments, the use of the I-SceI endonuclease was combined to a “COunter-selectable REporter” or CORE cassette in the frame of the *delitto-perfetto* technique, allowing the generation of a reciprocal translocation in a scar-less fashion [6]. Other techniques based on Cre/Lox recombination were used to make the genomes of *Saccharomyces cerevisiae* and *Saccharomyces mikatae* colinear and generated interspecific hybrids that produced a large proportion of viable but extensively aneuploid spores [7]. Cre/Lox recombination was also used to assess the impact of balanced rearrangements in vegetative growth and meiotic viability [8–10]. A novel approach using yeast strains with synthetic chromosomes allowed extensive genome reorganization through CreLox-mediated chromosome scrambling [11–14]. This approach proved to be efficient to generate strains with a wide variety of improved metabolic capacities [13,15–17]. Muramoto and collaborators recently developed a genome restructuring technology relying on a temperature-dependent endonuclease to conditionally introduce multiple rearrangements in the genome of *Arabidopsis thaliana* and *S. cerevisiae*, thus generating strains with marked phenotypes such as increased plant biomass or ethanol production from xylose [18]. Methods using Zinc Finger Nucleases (ZFNs) and Transcription Activator-Like Effector Nucleases (TALENs) were also developed to generate targeted rearrangement in yeast, mammalian and zebrafish cells [19–22]. Although these technologies provide very useful insights, they are often difficult to implement and/or rely on the use of genetic markers. For this reason, the development of the CRISPR/Cas9 (Clustered Regularly Interspaced Short Palindromic Repeats/CRISPR associated) system has boosted the field of genome engineering [23–25]. This system, initially derived from immune systems of bacteria, consists of an endonuclease encoded by the Cas9 gene of *Streptococcus pyogenes* and a short RNA that guides the endonuclease at the targeted genomic locus. The gRNA can be easily designed to target any genomic locus proximal to a “NGG” Promoter Adjacent Motif (PAM). This technology is now routinely used to introduce targeted DSBs in genomes from a wide variety of species [26]. In yeast, CRISPR/Cas9 induced DSBs can be repaired with high efficiency by providing homologous repair DNA cassettes, allowing a variety of genome editions. Previous studies achieved the introduction of point mutations, single and multiple gene deletions and multiplexed genome modifications at different loci by transforming cells with plasmids bearing single or multiple gRNAs and linear DNA repair templates [27–30]. CRISPR-based approaches have also been developed to add centromeres and telomeres to chromosome fragments [31] concatenating chromosomes [32,33] and for massively parallel genome editing to generate large libraries of genetic variants [34–36]. Interestingly, it has been noticed that multiplex genome editing can often result in undesirable chromosomal translocation suggesting that such rearrangements are likely programmable [37]. The ability to generate genome rearrangements with CRISPR/Cas9 has been known for some time [38] and more recently used to engineer translocations in mammalian cells with high efficiency. The principle was to introduce two DSB in two distinct chromosome with CRISPR, then repair the DNA ends in trans by HDR with donor DNA carrying a selection marker, lost in a second step by Cre/Lox recombination leaving a single loxP element at the chromosomal junction [39]. However, unequivocally determining the contribution of SVs to phenotype variations in a large variety of environmental conditions and independently from the contribution of single nucleotide variants cannot be envisioned in mammals because of overwhelming technical and ethical difficulties.

In this study, we developed two CRISPR-Cas9 genome editing strategies in yeast to generate markerless and scarless SVs with high efficiency in a control and unique genetic background, thus providing a mean to quantify their fitness impacts by high-throughput phenotyping independently from marker or background specific effects. The first strategy allowed generating on-demand any translocation at a base-pair resolution and we recapitulated the phenotypic consequences associated with known rearrangements. The second approach allows generating multiple SVs simultaneously leading to an important diversity of karyotypes. We showed that reshuffling the chromosome architecture between dispersed repeated sequences without disrupting any genes or changing their promoter sequences is sufficient to create fitness diversity in various stress conditions.

## RESULTS

### Rationale for chromosome reshuffling

We used a single-vector which encodes both the Cas9 nuclease gene and a gRNA expression cassette. This cassette allows cloning either a DNA fragment of 430 base-pairs reconstituting two different gRNAs in tandem or a unique 20 bp fragment corresponding to the target sequence of a single gRNA (fig 1A). This system is versatile as it allows generating either a single targeted or multiple translocations at once. Targeted translocations can be induced by a pair of gRNAs generating two concomitant DSBs in two different chromosomes. Any pair of gRNAs can be cloned in the vector in a single ligation step using as insert either a synthetic DNA fragment or a PCR product (see Methods). The two DSBs are repaired in trans upon homologous recombination with chimerical donor DNA resulting in targeted reciprocal translocations with a single-nucleotide precision (fig 1B). Multiple translocations are induced by generating several DSBs using a single gRNA that targets repeated sequences such as the Long Terminal Repeats (LTRs) scattered on different chromosomes. The multiple DSBs are repaired by recombination using the uncut copies of the LTRs as homologous templates (fig 1C). Both targeted and multiple translocations are engineered in a scar-less fashion and without integrating any genetic marker in the genome.

**Figure 1:**
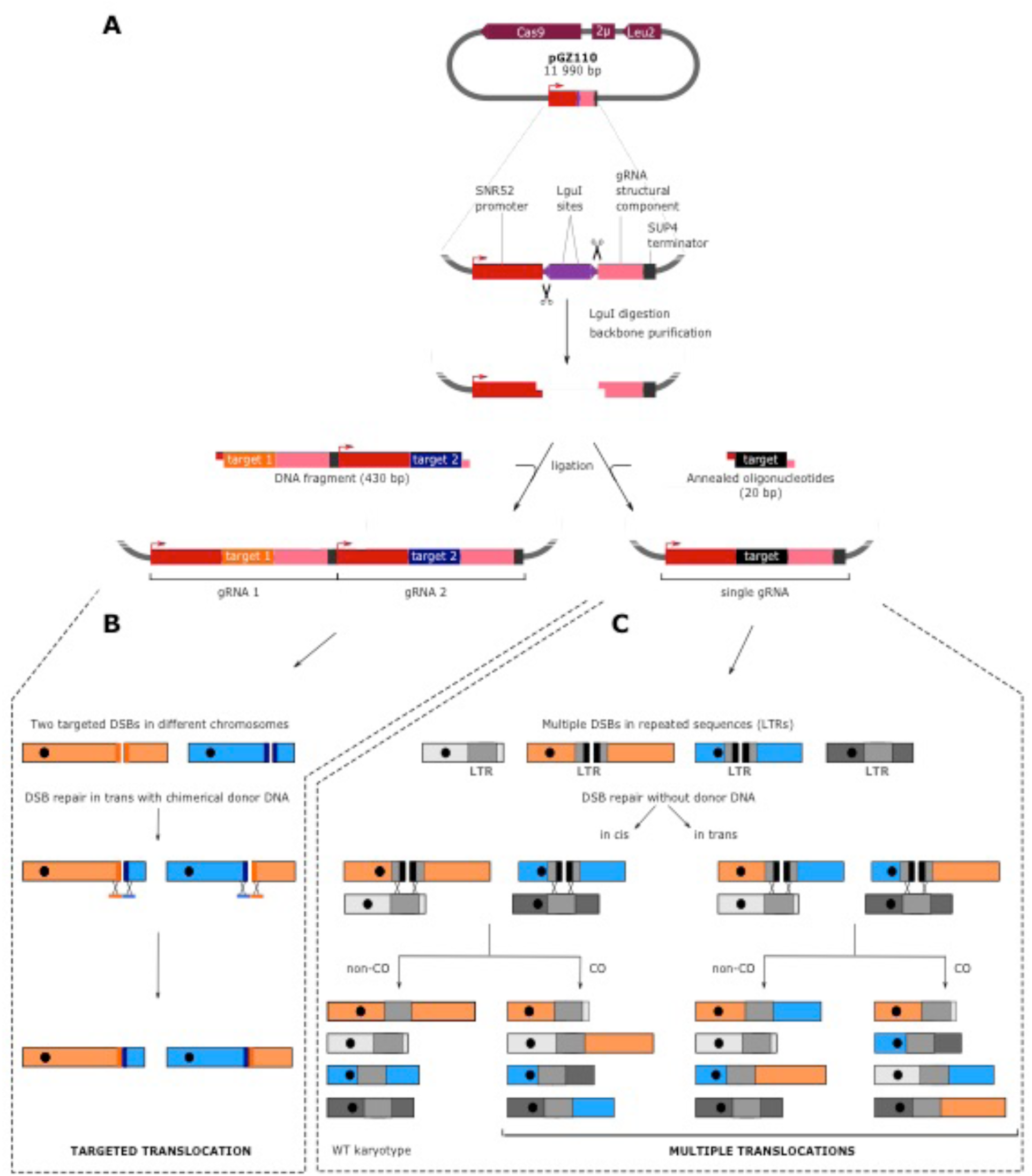
Strategy to reshuffle the yeast genome. **A.** Cloning gRNA target sequences in the pGZ110 vector (Bruce Futcher, Stony Brook University). Upon digestion, the two LguI sites generate non-complementary single strand overhangs of 3 bases. This system is versatile as it allows to clone in a single ligation step either a DNA fragment of 430 base-pairs reconstituting two different gRNA expression cassettes in tandem (left) or a 20 bp oligonucleotide corresponding to the target sequence of a unique gRNA (right). **B.** Induction of a targeted reciprocal translocation. Two DSBs are induced in different chromosomes and are repaired by homologous recombination with chimerical donor DNA oligonucleotides. The bold lines flanking the DSBs symbolize the target sequences. Donor DNAs force the repair to occur in trans (i.e. between two ends coming from the two different DSBs). **C.** Induction of multiple reciprocal translocations. This example illustrates a situation where the only two LTRs located in the blue and orange chromosomes are targeted by the gRNA (bold black lines indicate the target sequences) while LTRs located in the two grey chromosomes are not cut (devoid of target sequence). Note that more LTRs can be targeted as detailed below (see fig 5). DSBs are repaired by homologous recombination using the uncut LTRs as donor template. Repair can occur in cis (i.e. the two ends from the same DSB are repaired together) or in trans (i.e. two ends from two different DSBs are repaired together).

### Engineering a markerless reciprocal translocation at single base-pair resolution

We first engineered a reciprocal translocation between two reporter genes leading to phenotypes easy to observe upon disruption. Mutation in the *ADE2* gene involved in purine nucleotide biosynthesis results in the accumulation of a red pigment while mutating the *CAN1* gene which encodes an arginine permease confers canavanine resistance to the cells. To generate two concomitant DSBs on chromosomes V and XV carrying *CAN1* and *ADE2*, respectively, we cloned two previously described gRNA target sequences, namely CAN1.Y and ADE2.Y [27]. To repair the DSBs, we used donor DNA fragments of 90 base-pairs each composed of two homology regions of 45 bp identical to the sequences flanking CRISPR cutting sites (supp table 1). Two combinations of DNA repair donor fragments were used. As a control, we used donors called Point Mutation-donors (PM-donors), promoting the intra-chromosomal repair of DSBs in cis and mutating PAM sequences into stop codons, thus preventing further Cas9 activity. The donors promoting inter-chromosomal repair in trans, thus leading to a Reciprocal Translocation were called RT-donors. No point mutation needed to be introduced into the PAM sequence in this case because the translocation generates chimerical target sequences not complementary to the original gRNAs (fig 2A). Transformation with the plasmid bearing two gRNAs and either RT or PM-donors resulted in 95% and 76% of the colonies showing both the pink and resistance to canavanine phenotypes [ade2, can1], respectively (Material and Methods, fig 2B). In the RT-donor experiment the other 5% of colonies were all white and sensitive to canavanine, probably resulting from the transformation of the *LEU2* marker without induction of any DSB. In the PM experiment, we recovered 21% of white colonies, 83% of which being resistant to canavanine, likely resulting from a single DSB/repair in the *CAN1* gene. We also recovered 3% of pink and sensitive colonies likely resulting from a single DSB/repair in the *ADE2* gene (fig 2B). We then confirmed by PCR the presence of the chimerical chromosomal junctions of 16 [ade2, can1] strains recovered from the RT experiment. We further validated the translocation by karyotyping two [ade2, can1] strains by PFGE. No other visible chromosomal rearrangements could be observed apart from the expected translocated chromosomes VtXV and XVtV (fig 2C). Sanger sequencing of 250 bp around the chimerical junctions of these two strains confirmed that the translocation occurred right at the position defined by the sequence of the RT-donors with no additional mutations (supp fig 1).

**Figure 2:**
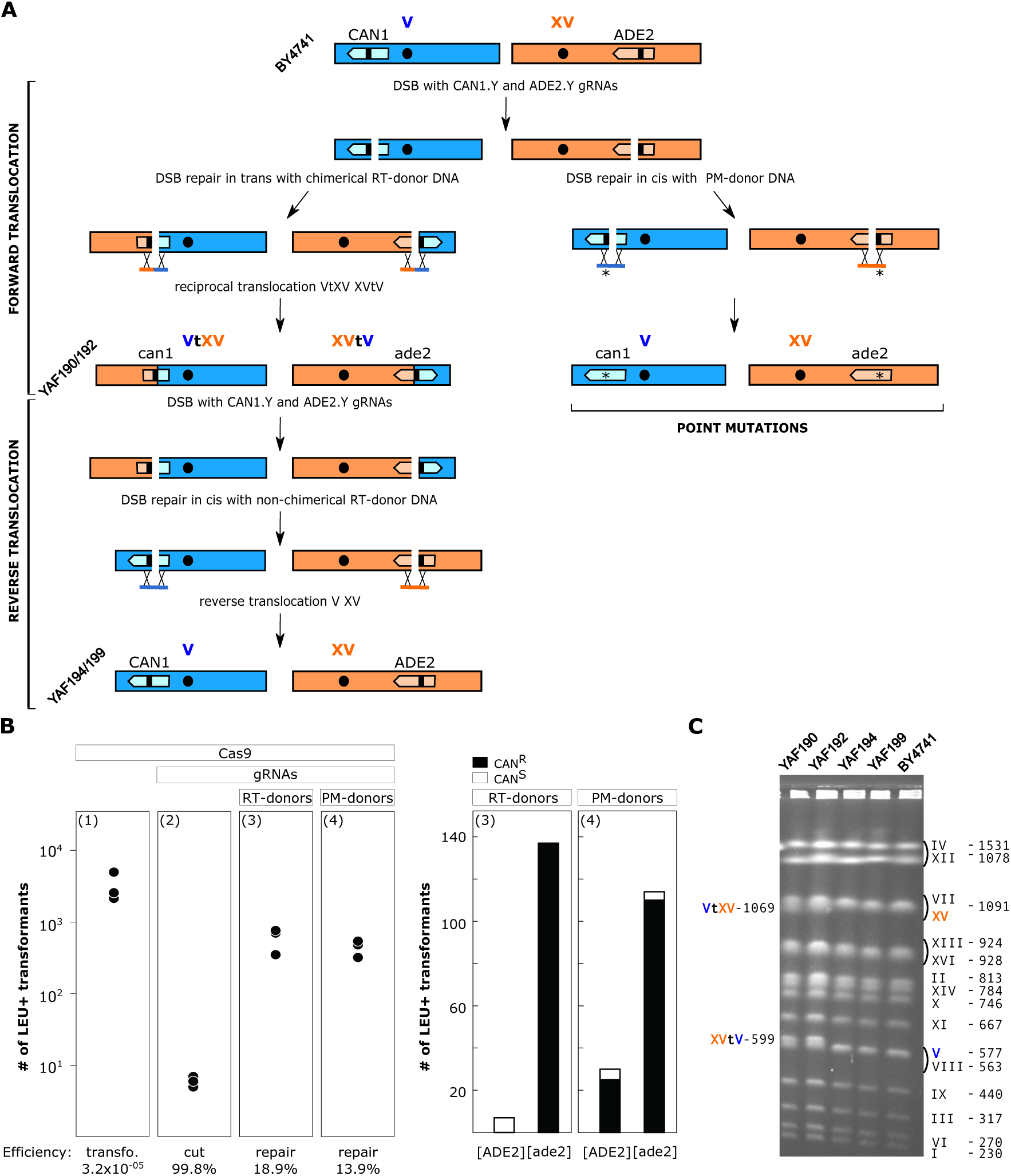
A Reversible markerless translocation. **A.** PAM sequences are symbolized by the small black bars within the *CAN1* and *ADE2* genes in dark blue and orange. RT-and PM-donors stand for Reciprocal Translocation and Point Mutation donor DNAs used to repair the DSBs. The point mutation responsible for the introduction of the STOP codon in the PAM sequence is indicated by an asterisk. Strain names are written in diagonal. **B.** Left: plots indicating the number of transformants for 10^8^ transformed cells obtained in 3 independent experiments. Panels (1) and (2) illustrate the efficiency of transformation with the Cas9 plasmid and cutting efficiency of the gRNAs, respectively. Panels (3) and (4) show DSB repair efficiency at the *ADE2* and *CAN1* loci by both the Reciprocal Translocation (RT) and Point Mutation (PM) donor DNAs (Methods). Right: Histograms of the numbers of white [ADE2], pink [ade2], canavanine resistant [CAN^R^] and sensitive [CAN^S^] colonies obtained with the RT (3) and PM donors (4), 100% and 96% of [ade2] transformants are also mutated for the *CAN1* gene, respectively. **C.** PFGE karyotypes of two strains carrying the *ADE2-CAN1* translocation (YAF190, YAF192) and two strains with the reverse translocations restoring the original chromosomes V and XV (YAF194 and YAF199 originating from YAF190 and YAF192, respectively). Chimerical chromosomes are denoted VtXV and XVtV. Original chromosome XV and V from the reference strain BY4741 are indicated in orange and blue respectively. Chromosome size is indicated in kb.

We next showed that our system allows to sequentially generate several targeted translocations. To illustrate this we performed the reverse translocation to restore the WT chromosomes V and XV and functional *ADE2* and *CAN1* genes in the strains that carried the *ADE2*-*CAN1* translocation (YAF190 and YAF192 in fig 2C). We took advantage of the high instability of the Cas9 plasmid that can be easily cured from the strains (Material and Methods). We first cured the plasmid from the two translocated strains. Then, we cloned a pair of gRNAs that target the chimerical junctions formed by the *ADE2*-*CAN1* translocation in the Cas9 plasmid and designed repair donors to restore *ADE2* and *CAN1* to their original configuration (fig 2A, supp table 1). Co-transformation with the gRNAs plasmid and donor fragments resulted in 94.2% of colonies which restored the white and canavanine sensitive phenotypes [ADE2, CAN1]. We performed pulse field gel electrophoresis (PFGE) karyotyping of two [ADE2, CAN1] strains. No difference could be observed between the karyotype of these two strains and that of the original BY4741 strain (YAF194 and YAF199 in fig 2C). The chromosomal junctions of the two de-translocated strains were Sanger-sequenced and were found identical to BY4741 natural junctions (supp fig 1). These results demonstrate that sequential chromosomal translocations can be engineered at base-pair resolution with high efficiency.

Finally, we also showed that our system can be used to simultaneously generate a deletion of a few nucleotides and a reciprocal translocation. We used the same CAN1.Y and ADE2.Y gRNA target sequences as above and designed new donor fragments inducing deletions of 27 and 23 bp, including the PAM sequences, on chromosome V and XV, respectively (supp fig 2A supp table 1). As above, we obtained a high proportion of [*ade2, can1*] transformants (96%). PCR of the junctions and karyotyping of 4 strains showed that chromosome V and XV underwent the expected reciprocal translocation (fig 3 A). The genome of one translocated strain was Nanopore sequenced and *de-novo* assembled (Material and Methods, supp table 2 and 3) to check whether off-target activity of the Cas 9 nuclease could result in unexpected additional rearrangements. The translocated and reference genomes were entirely collinear except for the expected translocation and associated deletions on chromosomes V and XV (fig. 3B). No other rearrangement was observed, suggesting no major off-target activity of the Cas9 nuclease in this strain. In addition, the junction sequences are identical to the sequences of the chimerical donor fragments (supp fig 2B). These experiments demonstrate that a deletion and reciprocal translocation can be concomitantly engineered at base-pair resolution with CRISPR/Cas9 in the yeast genome.

**Figure 3:**
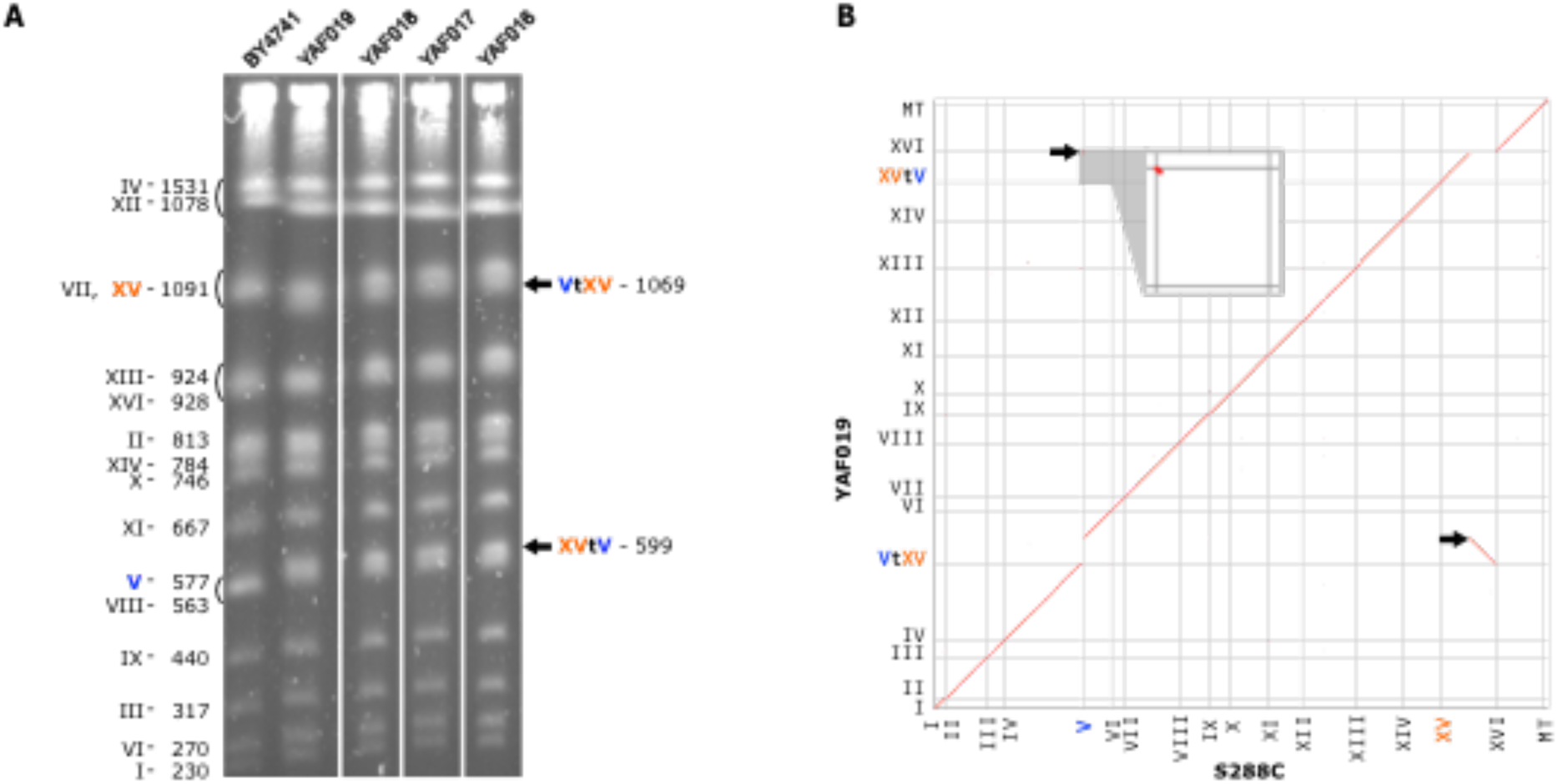
A non-reversible translocation between *ADE2* and *CAN1* genes. (A) PFGE karyotypes of 4 independent strains carrying the translocation (YAF016 to YAF019). All symbols are identical to Fig. 2C. (B) Homology matrix of *de-novo* assembled strain YAF019 vs S288c reference genome. Translocated fragments are indicated by black arrows.

### Recapitulating a natural translocation involved in sulphite resistance in wine strains

It was previously reported that a reciprocal translocation between the promoters of *ECM34* and *SSU1*, a sulphite resistance gene, created a chimerical *SSU1-R* allele with enhanced expression resulting in increased resistance to sulphite in the wine strain Y9J [40]. This translocation resulted from a recombination event between 4 base-pair micro-homology regions on chromosomes VIII and XVI. We therefore engineered the same translocation into the BY4741 background. We designed two gRNA target sequences as close as possible to the micro-homology regions (supp fig 3A and supp table 1). The first gRNA targeted the *SSU1* promoter region, 115 base-pairs upstream of the start codon. The second gRNA targeted the promoter region of *ECM34*, 24 base-pairs upstream of the start codon. To mimic the translocated junctions present in the wine strains, we designed 2 double stranded synthetic DNA donors of 90 base-pairs centered on the micro-homology regions but not on the cutting sites. In addition, each donor also contained a point mutation in the PAM sequences to prevent subsequent CRISPR recognition (supp fig 3A).

The transformations with the Cas9 plasmid containing the two gRNAs and the donor DNA yielded on average 202 transformants. We tested natural and chimerical junction by colony PCR for 16 transformants and found the expected chimerical junctions in 15 of them. This translocation was not visible by PFGE karyotyping because the size of the translocated chromosomes were too close to the size of the original chromosomes (XVI 948 kb, VIII 563 kb, VIIItXVI 921 kb and XVItVIII 599 kb). To further validate the rearrangement, we checked the junctions by southern blot for one translocated strain with probes flanking the two cutting sites on chromosome VIII and XVI (supp fig 3B). This experiment validates the presence of the chimerical junction fragments in the rearranged strain YAF082 as compared to the WT. Finally, we sequenced the junctions of the translocated strain and found that the rearrangement occurred within the expected micro-homology regions (supp fig 3C). In addition, the system was designed such that both mutated PAMs ended up into the promoter of *ECM34* the translocation, therefore unlikely to have any impact on the expression of the sulphite resistance gene *SSU1* (supp fig 3C).

We then compared sulphite resistance between the lab strain in which we engineered the translocation (YAF202 corresponding to YAF082 cured for the Cas9 plasmid, see Methods), the non-translocated parental lab strain (BY4741) and wine isolates that carry or not the translocation of interest (Y9J and DBVPG6765, respectively). Surprisingly, we found that the engineered strain with the translocation was the least resistant of all strains with a Minimal Inhibitory Concentration (MIC) of 1 mM (fig. 4). By comparison, the reference strain and the wine isolate without the translocation both had a MIC of 2 mM. This suggests that the promoter of *ECM34* is weaker than the *SSU1* promoter in the BY background. As expected, the wine isolate with the translocation (Y9J) was the most resistant of all (MIC > 20 mM, fig. 4), However, it was reported that this wine strain also had four tandem repeats of a 76 bp segment in the promoter region of *SSU1* originating from the *ECM34* locus and it was shown that the number of repeats positively correlated with sulphite resistance [40]. This suggested that the translocation would not be *per se* responsible for increased resistance. Resistance would in fact result from the repeats in the *ECM34* promoter region that were brought in front of the *SSU1* gene by the translocation. There is only one copy of this 76 bp sequence in the BY background. To test whether promoter repeats were responsible for the resistance phenotype we PCR amplified the repeat-containing promoter of the translocated wine strain Y9J (introducing a point mutation in the PAM sequences to avoid subsequent recognition by Cas9, supp table 1) and used the PCR product as donor in a CRISPR experiment to introduce the promoter repeats in front of the *SSU1* gene in the translocated BY4741 strain. We designed a gRNA targeting the region between the single copy motif and the beginning of *SSU1* (supp table 1). We obtained hundreds of transformants and found that 5 out of 8 transformants tested by PCR contained the four tandem repeats and were subsequently validated by Sanger sequencing (supp fig 3D). The addition of the repeats in the *SSU1* promoter in the translocated lab strain (YAF158) resulted in increased sulphite resistance with a MIC of 7 mM (fig. 4) therefore confirming the effect of the repeats on the resistance phenotype. However, the chromosomal configuration and the promoter repeats are the same in YAF158 and the Y9J wine strain, yet the wine strain is much more resistant than the lab strain suggesting that additional polymorphisms must contribute to the phenotype in the wine isolate.

**Figure 4:**
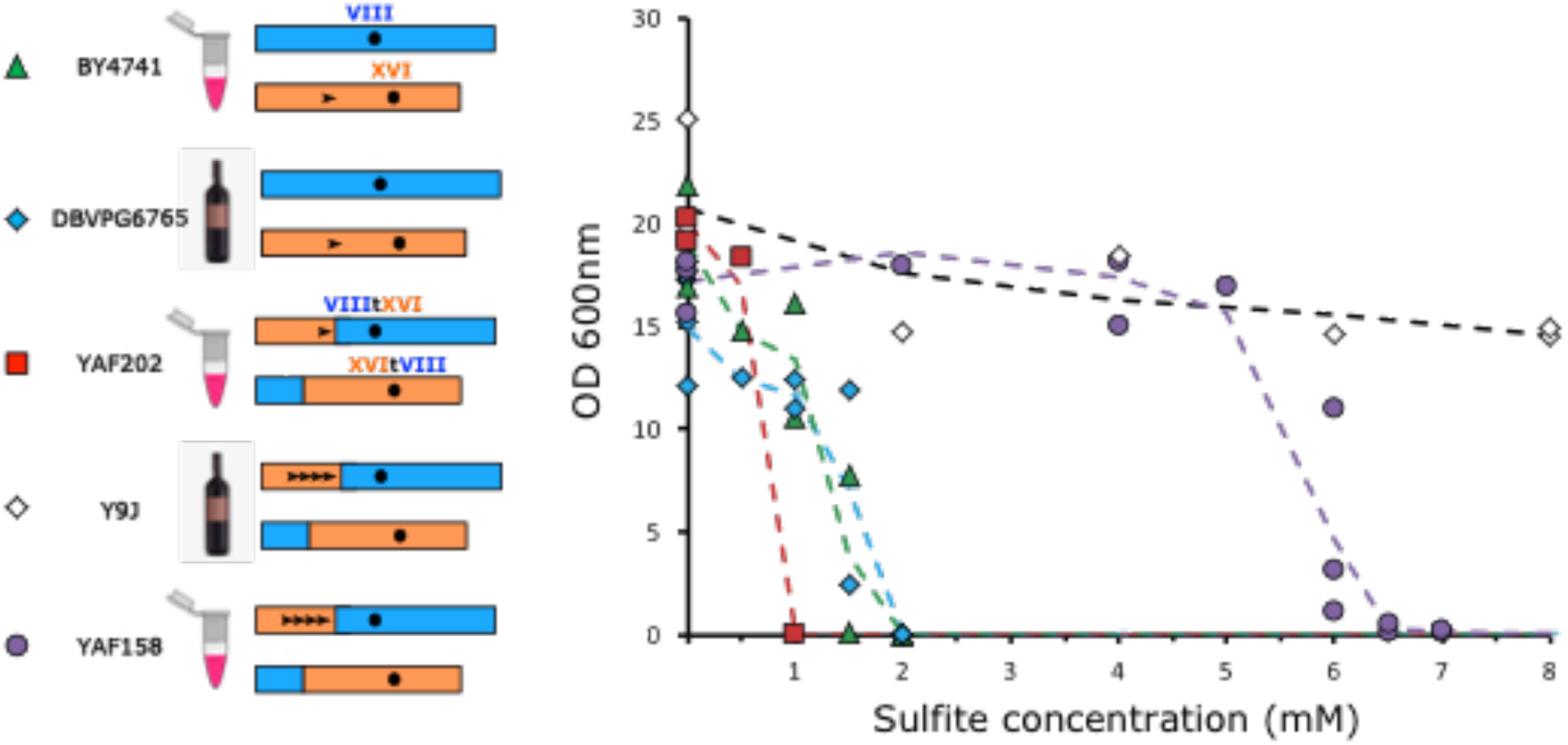
Quantification of sulphite resistance. Laboratory and wine strains are symbolized by eppendorf tubes and bottles, respectively. The chromosomal configurations are represented by blue and orange rectangles. Black arrowheads symbolize the 76 bp repeats in promoter regions. The determination of the MIC of sulphite was performed for the entire range of concentration for all strains but for readability all points with a null value of OD for concentrations higher than the MIC were omitted from the graph.

### Reshuffling chromosomes with multiple translocations

It has been previously shown that DSBs give rise to chromosomal rearrangements when they fall in dispersed repetitive elements such as Ty retrotransposons [1]. We reasoned that generating in a single step multiple DSBs targeting Ty repeated sequences should also result in genome reshuffling through multiple translocations without altering any coding sequence or promoter region. We chose a gRNA sequence flanked by a PAM that targets 5 different Ty3 LTR copies located in chromosomes IV, VII, XV and XVI (four of which comprise a region identical to the gRNA target sequence while the fifth copy differs from the target sequence by a single mismatch at its 5’ end, fig 5A). In addition, there are 30 other complete copies of Ty3 LTRs dispersed throughout the genome but they contain several mismatches or indels relatively to the sequence of the gRNA and/or are devoid of PAM, suggesting that Cas9 will not cut at these sites (fig 5A). We hypothesized that these uncut LTRs would be used as internal donor templates for DSB repair. We chose to target these five Ty3 LTRs because inducing DSB at these locations should allow significant genome reshuffling without compromising to much the viability of the cells by overloading the repair system by too many DSBs. DNA ends originating from the five DSBs can be repaired both in cis, *i.e.* the two ends from the same DSB are repaired together or in trans, *i.e.* the two ends from two different DSBs are repaired together. A WT-like karyotype is expected when all DSBs are repaired in cis and without any crossover with the internal Ty3 donor templates (type A in fig 5B). In addition, we can predict 23 viable combinations of rearranged karyotypes when all DSBs are repaired in trans without crossover (types B to X in fig 5B). Note that only reciprocal translocations are expected because the two targeted Ty3 LTRs on chromosome XV are in the same transcriptional orientation and therefore cannot induce the inversion of the intervening segment. In addition, all rearranged karyotypes comprising acentric or dicentric chromosomes are supposedly non-viable and thus will not be recovered. Other combinations of viable karyotypes resulting from both cis and trans repair with crossovers within uncut LTR donors could also be generated but are hard to predict because of the large number of possible LTR donors that can be used as template for repair and therefore are classified as ‘unpredicted’ in the following.

**Figure 5:**
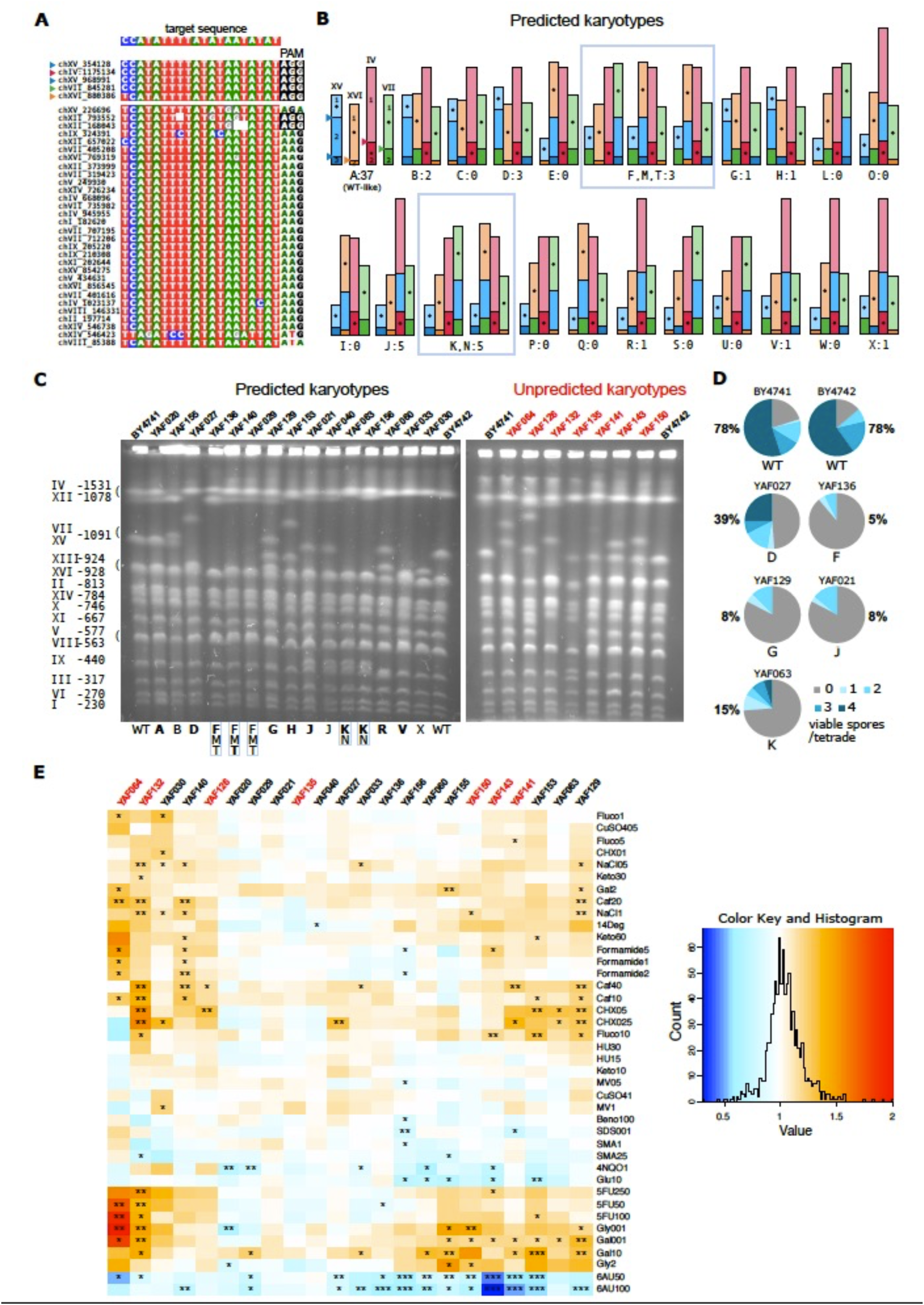
Induction of multiple rearrangements and phenotypic diversity of reshuffled strains. **A.** Multiple alignment of the target regions inTy3-LTR sequences. The gRNA target sequence is indicated on the top and the corresponding homologous regions in the LTR are highlighted below. Associated PAM sequences are highlighted in black. The five LTR sequences at the top are the best matches to the gRNA target sequence. Coordinates on the left indicate the start of the corresponding TY3 LTRs. **B.** Predicted rearranged karyotypes (types A to X). Chromosomes are represented proportionally to their size in kb. Centromeres are represented by black dots located in the middle of their carrying fragment for readability. The chromosomal location of the 5 cutting sites corresponding to the 5 best matches to the gRNA are indicated by colored triangles on the type A profile. The number of strains of each type that were characterized by PFGE is indicated below each drawing. Types B to F have only 2 chimerical junctions resulting from a single reciprocal translocation between 2 chromosomes. Types G to M have 3 chimerical junctions resulting from translocations between 3 chromosomes (G, H, L, M) or the transposition of the chromosomal fragment XV.2 (I, J, K). Types N to V have 4 chimerical junctions resulting from a combination of translocations and transpositions. Types W and X have all 5 chimerical junctions. **C.** PFGE of 23 strains with predicted (left) and unpredicted (right) karyotypes. The control WT strains (BY4741 and BY4742) are located on the most external lanes of each gel and their chromosome size is indicated on the left. Strain names and types are indicated above and below the gels, respectively. The predicted types are in bold when all chimerical junctions were validated by PCR. **D.** Percentage of viable spores and proportion of tetrads with 0, 1, 2, 3, 4 viable spores obtained from crosses between BY (WT and rearranged) and WT SK1 strains. **E.** Phenotypic variation among reshuffled strains. The heatmap represents the growth ratio of each strain (*i.e.* the colony size on the tested conditions divided by its size on SC) divided by the growth ration of BY4741 or BY4742, depending on the origin of the shuffled strain (Methods). The stars indicate the significant phenotypic effects (*, ** and *** indicate pval<10^-2^, 10^-3^ and 10^-4^, respectively). Refer to supplementary fig 4 for the list of abbreviations corresponding to the 40 growth conditions. The strain names in red correspond to the unpredicted karyotypes in panel C.

We transformed BY4741 and BY4742 cells with the Cas9/gRNA plasmid targeting the 5 Ty3 LTRs and recovered a total of 211 and 159 transformants, respectively. We PFGE karyotyped 69 transformants (37 BY4741 and 32 BY4742) out of which 30 showed clear chromosomal rearrangements on the gels, representing 18 different karyotypes in total (fig 5C). This result demonstrates that genomes are efficiently reshuffled via our strategy. In total 23 strains showed predicted rearrangements, representing 10 distinguishable profiles (B; D; (F,M,T); G; H; J; (K,N); R; V and X in fig 5B and 5C). We validated the presence of most predicted junctions by colony-PCR in 18 out of the 23 strains (fig 5C). We sequenced all chromosomal junctions in 2 strains that show the most frequently observed rearranged profile (type J in fig 5B and 5C) and found that all junctions, from both chimerical and un-rearranged chromosomes, were mutated in their PAM compared to the original target sequence (supp fig 4). This shows that during shuffling, all targeted sites were cut and repaired using as donor uncut Ty3 LTRs that had no PAM. Additional mutations in the region corresponding to the gRNA target sequence were also observed for 3 junctions, but were too few to identify which copy of the uncut Ty3 LTR was used as donor (supp fig 4). We tested any possible over-or under-representation of the predicted rearranged types by randomly sampling with replacement 15 draws (corresponding to the 15 characterized strains) in a uniform distribution of 18 types (24 types excluding type A (not rearranged) and types (F,M,T) and (K,N) that are not discernible). We performed 10,000 realizations and counted the number of times we sampled the different types. The probability to observe 5 times the predicted type J was 0.017 which suggests that this type could be overrepresented. No other type showed deviation from expectation. However, given the relatively small number of characterized strains, we cannot exclude that DSBs would be repaired in a random way.

Moreover, we obtained 7 strains with distinct unpredicted karyotypes involving chromosomes other than the 4 targeted ones (fig 5C). For instance, chromosomes XI and XIV that have no PAM sequence associated with their Ty3 LTRs (fig 5A) are absent from several karyotypes showing that they can be rearranged in the absence of DSB (fig 5C). Surprisingly, some of karyotypes showed an apparent genome size increase suggesting the presence of unexpected large duplications (see YAF064, YAF126, YAF143 and YAF150 in fig 5C). Using Oxford Nanopore MinION, we sequenced and *de-novo* assembled the genome of YAF064. We characterized in this strain an unequal reciprocal translocation between chromosomes VII and XV (supp fig 5A). The junctions corresponded to the targeted CRISPR cut site on chromosome XV but not on chromosome VII. In the chimerical chromosome XVtVII, the junction occurred away from the expected site resulting in a 30 kb increase in DNA content. In the chimerical chromosome VIItXV, the junction also occurred away from the expected site but was accompanied by a truncated triplication of a 110 kb region. The missing part of the triplication corresponds to the 30 kb region found in the reciprocal chromosome XVtVII. The sequencing coverage relatively to the reference genome clearly confirmed the triplication of the complete 110 kb region (supp fig 5B). Interestingly the 110 kb segment, composed of regions *a* and *b*, is flanked by two Ty3 LTRs. Moreover, two full length Ty2 retrotransposons are found, one in chromosome XV directly flanking the targeted Ty3 LTR and the other one in chromosome VII at the junction between regions *a* and *b* (supp fig 5C).

Finally, we also recovered 37 type A strains that had a WT karyotype (fig 5B and C). For all of them we validated the presence of all un-rearranged junctions by PCR. We sequenced the junctions in 4 independent strains. We found that these strains underwent two different paths within the same experiment. Firstly, 3 clones had all junctions identical to the original LTR sequences with intact PAM, suggesting that Cas9 did not cut the target sites. Secondly, in the fourth clone, the PAM sequences of three sequenced junctions were mutated, showing that the corresponding chromosomes were cut and repaired yet without any rearrangement (supp fig 4). In this clone, one junction could not be amplified, possibly because of a small-sized indel that could not be observed on the PFGE profile.

### Exploring the phenotypic diversity of reshuffled strains

It is well known that strains bearing heterozygous translocations have impaired meiosis resulting in low spore viability [8,41–43]. We checked that crosses between rearranged and parental strains indeed produced few viable spores (fig 5D). We tested 5 different strains with predicted rearranged karyotypes and PCR-validated junctions. These strains were chosen because they encompassed various rearrangements including reciprocal translocations between 2 and 3 chromosomes (types D, F and G, respectively) and transpositions (types J and K). For each cross, we dissected 10 tetrads from two independent diploid strains (20 tetrads in total). The control strain without any rearrangement shows 78% of viable spores with most tetrads harbouring 3 to 4 viable spores (fig 5D). By contrast, all heterozygous diploids had a severely impaired spore viability ranging from 5 to 39% with predominantly only 1 to 2 viable spores per tetrad. We observed no clear correlation between the type of rearrangement and the impact on fertility. One strain carrying a single reciprocal translocation between 2 chromosomes (type F) showed the lowest viability, comparable to that of a more rearranged strain with 2 translocations between 3 chromosomes (type G). These results show that knowing the type of rearrangements is not sufficient to predict their quantitative impact on meiotic fertility, although most of them have a drastic negative effect.

Secondly, 22 rearranged strains (the 15 predicted and 7 unpredicted karyotypes presented on fig 5C) were phenotyped in 40 different growth conditions impacting various physiological processes and in complete synthetic media as the reference condition. These 40 conditions include various types of stresses, different drugs interfering with replication, transcription and translation as well as compounds impacting several subcellular structures (supp table 4). In total, we performed 943 phenotypic measurements (fig 5E). We identified about twice as many cases where reshuffled strains grew significantly faster compared to cases where they grew slower than the WT strain (91 and 48 cases, respectively), suggesting that genome shuffling might be advantageous in many stressful conditions. The strongest phenotypic advantages of all corresponded to the strain with the unpredicted karyotype harboring the 110 kb triplication (YAF064, see above) when DNA synthesis is impaired (in the presence of the pyrimidine analog 5-fluorouracile) and in starvation (low carbon concentration 0.01% of galactose or glycerol, fig 5E). However, none of 36 triplicated genes with a known function is directly involved either in DNA synthesis or starvation, suggesting that the other 18 dubious ORFs present in this region could be involved in these phenotypes. More generally, for the conditions that produce the greatest effects, most of the strains tended to react in a similar way. For instance, in the presence of 6-azauracil (6AU), 4NQO and high glucose concentration all the strains that showed a significant phenotypic variation grew slower than the WT while in the presence of galactose, caffeine, cycloheximide and fluconazole all the strains that showed a significant variation grew faster than the WT (fig 5E). Most strains (19 out of 22) showed variations in at least 2 different conditions showing that genome shuffling is efficiently broadening the phenotypic diversity. The most variable strain, YAF132, presented significant growth variations in 17 out of the 40 conditions (faster and slower than the WT in 15 and 2 conditions, respectively). By opposition, the 2 type J strains (YAF021 and YAF040) as well as one strain with an unpredicted karyotype (YAF135) showed no phenotypic variation in nearly all the 40 conditions. In conclusion, these experiments revealed that reorganizing gene order along the chromosomes has direct phenotypic consequences, including fitness advantage in many environmental conditions.

## DISCUSSION

There are mounting evidences that SVs play a major role in phenotypic variation [44–46]. However, these genetic variants are the most difficult to interpret with respect to their functional consequences. In this study, we developed a versatile CRISPR/Cas9-based method allowing to engineer, with the same efficiency as point mutations, both uniquely targeted and multiple reciprocal translocations. Our method therefore provides the possibility to quantify the role played by structural variants in phenotypic diversity in a wide range of environmental conditions and in any genetic background.

Targeted reciprocal translocations were obtained with a single-nucleotide precision by cloning pairs of gRNAs that induce two concomitant DSBs in different chromosomes, which were repaired by chimerical donor DNA (see fig 1A and B). After the translocation, the new chromosomal junctions are not recognized by Cas9 which makes the mutation of the PAM sequence unnecessary. The Cas9 plasmid is highly unstable and therefore can easily be cured from the strain which allows to perform successive rounds of transformation to sequentially accumulate several targeted translocations in the same genome. In addition, our methods also allow to introduce deletions at the breakpoint concomitantly with the induction of the translocation.

The versatility of our approach allows to untangle the phenotypic impact of SVs from that of the genetic background. We illustrated this with the translocation between the *SSU1* and *ECM34* genes providing increased sulphite resistance to wine isolates, supposedly as a consequence of sulphite supplementation in wine-making [40]. Engineering the same translocation in the BY4741 background resulted in decreased sulphite resistance. Therefore, the same translocation produces two opposite phenotypes in the laboratory and the wine backgrounds. It was previously reported that the two strains also differ by a 76 bp repeated motif in the promoter of *ECM34* and that sulphite resistance correlated with the number of repeats [40]. Adding the four copies of the wine promoter into the translocated BY4741 strain that is devoid of repeats resulted in increased sulphite resistance, confirming that the repeats rather than the translocation are responsible for sulphite resistance. However, sulphite resistance remained lower in the laboratory as compared to the wine isolate despite they share both the same translocation and promoter repeats. This strongly suggests that additional polymorphisms such as SNPs also contribute to sulphite resistance in the wine isolate. For instance, there are 4 non-synonymous SNPs in the coding sequence of *SSU1* between the 2 genetic backgrounds. These findings provide a striking example of the advantages brought by our technique to untangle the phenotypic impact of SVs from that of the genetic background. Multiple translocations were obtained by using a single gRNA that targets dispersed repeat sequences leading to a large diversity of karyotypes. These results show that the genotypic space accessible by such an approach is probably very large. Our genome reshuffling procedure is reminiscent of the restructuring technology developed by Muramoto and collaborators [47] that uses a temperature-dependent endonuclease to conditionally induce multiple DSBs in the genome of yeast and *A. thaliana*. However, our method is potentially more versatile because we can target different types of repeated sequences with variable locations (within or between genes, subtelomeric or internal, etc) and variable copy numbers. By analyzing the genome of *S. cerevisiae* for repeated gRNA target sequences, we identified 39,374 repeated targets with varying number of occurrences in the genome (from 2 to 59). Repeated targets were mostly found within Ty transposable elements (11,503) and protein coding genes (10,368 in 479 genes located in all 16 chromosomes). Interestingly, we also found 3,438 repeated target gRNA located in intergenic regions which should allow to rearrange chromosomes without disrupting coding sequences. In addition, our CRISPR/Cas9 shuffling procedure is doable in any genetic background and therefore potentially more versatile than the Cre/Lox based SCRaMbLE techniques that can be used only in yeast strains with synthetic chromosomes.

Given the location of the multiple DSBs induced by the Ty3 LTR gRNA we chose and considering that all viable rearranged chromosomes must have a single centromere, we could predict the formation of 23 rearranged karyotypes (types B to X in fig 5B). These translocations result from trans repair between targeted DSBs (*i.e.* two DNA ends from two different DSBs are repaired together) without any crossover within the uncut LTRs used as template (fig 1C). We isolated 23 strains representing 10 different predicted karyotypes (fig 5C). The observed distribution of all predicted types, except type J, was very similar to the expected frequency of each types, suggesting that DSBs were repaired in a random way and that no specific chromosomal contact would be favored during repair. It is interesting to note that type J corresponds to the only chromosomal combination amongst all discernible predicted types where chromosome XVI remains non-rearranged (fig 5B). The target sequence on chromosome XVI is the only one that contains a mismatch with the gRNA sequence (fig 5A), suggesting that even a single mismatch located on the 5’ end of the target sequence could be sufficient to decrease the cutting efficiency contrarily to *in vitro* results showing that mismatches at the last two nucleotides at the PAM distal region show similar or even higher cleavage efficiency than that of WT target [48,49]. We also isolated 7 strains with rearranged karyotypes different from the 23 predicted profiles (fig 5C). Such rearrangements are believed to result from crossovers involving uncut LTRs (fig 1C), leading to translocations involving uncut chromosomes. The large number of LTRs that can be used for each repair event explains why these karyotypes would be difficult to predict. Moreover, we cannot exclude that some of the unpredicted karyotypes would also result from untargeted LTRs being cut by CRISPR/Cas9. However, this scenario seems very unlikely because all other LTRs are devoid of PAM and/or contain multiple mutations and indels in the target sequence (fig 5A). Another possible explanation to the presence of unpredicted karyotypes would be multipartite ectopic recombination between cut and uncut chromosomes where a single end of a DSB can invade multiple sequences located on intact chromosomes during its search for homology leading to translocations between uncut chromosomes by multi-invasion-induced rearrangement [50]. Surprisingly, more complex rearrangements involving large duplications were also recovered (fig 5C, supp fig 5), showing that inducing DSB can trigger the formation of large duplicated segments, as previously described [51]. Consistently, it has been shown that BIR can occur by several rounds of strand invasion, DNA synthesis and dissociation within dispersed repeated sequences such as LTRs, leading to chromosome rearrangements [52]. One such example was found here in the strain with the large segmental triplication where one chimerical junction probably resulted from template switching between the Ty3 LTR and the full-length Ty2 elements found in the vicinity of the CRISPR-induced DSBs (supp fig 5).

We determined the phenotypes of strains carrying multiple rearrangements both in mitotic and meiotic conditions. We showed that crosses between rearranged and parental strains produced very few viable spores (fig 5D), supporting earlier studies revealing that meiotic pairing and segregation is impaired in diploid strains bearing heterozygous translocations [8,41–43]. We also showed that genome reshuffling generates a great phenotypic diversity under specific environmental conditions including many cases of fitness advantage (fig 5E). Similar findings were previously described both in *S. cerevisiae* [53,54] and *Schizosaccharomyces pombe* [8,46]. However, for the first time in this study all rearrangements are completely markerless and scarless (no CRE/Lox site) and for the predicted karyotypes no gene was disrupted nor duplicated suggesting that balanced rearrangements between repeated sequences such as Ty3 LTRs, simply reconfiguring the chromosome architecture are sufficient to create fitness diversity. Ty3 LTRs control expression of the Ty3 genomic RNA and have an organization similar to other yeast promoters. Ty3 LTRs contain positive control elements for pheromone induction, negative control elements for mating-type and also pheromone-independent transcriptional activity [55]. The transcriptional activity of Ty3 LTRs could possibly explain the observed phenotypic diversity associated with genome reshuffling at these sites. Interestingly, the presence of the nucleotide-depleting drug 6AU induces the strongest growth defect of all tested conditions (fig 5E). Sensitivity to 6AU is a well-documented phenotype associated with transcription-elongation mutants, reducing both the elongation rate and processivity [56–59], suggesting that transcription might be globally affected in the reshuffled strains. Further work is needed to decipher the molecular mechanisms at the origin of the phenotypic diversity associated with genome reshuffling.

## METHODS

### Strains and media

The strains of *Saccharomyces cerevisiae* BY4741, (*MAT*a, *his3Δ1, leu2Δ0, ura3Δ0, met15Δ0*) and BY4742 (*MAT*α, *his3Δ1, leu2Δ0, ura3Δ0, lys2Δ0*) were used for generating both targeted and multiple translocations. Pre-cultures were performed in YPD (yeast extract 10 g.l^-1^ peptone 20 g.l^-1^, glucose 20 g.l^-1^) or in SC (Yeast Nitrogen Base with ammonium sulfate 6.7 g.l^-1^, amino acid mixture 2 g.l^-1^, glucose 20 g.l^-1^) for high-throughput phenotyping experiments. After transformation, cells were selected on SC medium depleted in leucine. The SK1 strains SKY1513 (*MAT*α, *ho::LYS2, ura3, leu2::HISG, lys2, arg4*(Nnde1)-Nsp, *thr1-A, SPO11*-HA3-His6::*KanMX4*) and SKY1708 (*MAT*a, *ho::LYS*2, *ura3, leu2::HISG, lys2, arg4*-Bgl::NdeI-site1°, *CEN8::URA3*) were used for their high sporulation efficiency compared to BY strains [60] to perform crosses and to quantify spore viability of the rearranged strains. Sulphite resistance of our engineered strains (YAF202 and YAF158) was compared in liquid assay to that of wine strains DBVPG6765 (provided by Gianni Liti (IRCAN, Nice)) and Y9J_1b and to the BY4741 strain. Cells were grown overnight in YPD broth prior to inoculation at a concentration of 10^4^ cells/mL in 8ml of YPD broth buffered at pH=4 with tartaric acid and containing Na2SO3 concentrations ranging from 0 to 20 mM [61]. The 50 ml culture tubes were tightly closed to avoid evaporation of sulphites and incubated at 30°C. Photographs and OD600 were taken after 48 hours to quantify cell growth. Plasmid cloning steps were performed in chemically competent *Escherichia coli* DH5α. Ampicillin resistant bacteria were selected on LB medium supplemented with ampicillin at 100 µg/ml.

### Identification of CRISPR/Cas9 target sequences

For the translocation between *ADE2* and *CAN1*, the target sequences CAN.Y and ADE2.Y found in the literature [27] were re-used. The new targets formed by the first translocation were targeted to “reverse” the translocation and restore the original junctions. For the *ECM34*/*SSU1* translocation, specific CRISPR/Cas9 target sequences with minimal off-targets were chosen as close as possible to the natural recombination site of wine strains with the CRISPOR v4.3 website (http://crispor.tefor.net) using the reference genome of *Saccharomyces cerevisiae* (UCSC Apr. 2011 SacCer_Apr2011/sacCer3) and the NGG protospacer adjacent motif. For multiple rearrangements, we identified 39 occurrences of Ty3-LTRs in the latest version of the genome of *Saccharomyces cerevisiae* S288C (accession number GCF_000146045.2), which we have aligned using MUSCLE. Four sequences were incomplete and excluded from further analysis. We then manually selected a suitable gRNA sequence targeting five Ty3-LTR elements and looked for off-targets to this target sequence with CRISPOR. Predicted off-targets had either mismatches with the chosen guide and/or were devoid of PAM indicating that they would not be recognized by CRISPR/Cas9.

### Construction of CRISPR/Cas9 plasmids with one or two guides

The original plasmid pGZ110 was kindly provided by Bruce Futcher (Gang Zhao, Justin Gardin, Yuping Chen, and Bruce Futcher, Stony Brook University; personal communication). All plasmids constructed in this study were obtained by cloning in pGZ110 either a 430 bp DNA fragment reconstituting a two-gRNA expression cassette or a 20 bp fragment corresponding to the target sequence of a single gRNA (fig 1a). We linearised pGZ110 with the enzyme LguI (ThermoFischer FD1934) and gel purified the backbone. In order to clone a single 20 bp target sequence we first annealed two oligonucleotides of 23 bases with 5’ overhangs of 3 bases complementary to the LguI sites. Annealing was performed by mixing equimolar amounts of forward and reverse oligonucleotides at 100 pmol/µl with NEBuffer 4 (New England Biolabs), heating 5 minutes at 95°C and allowing the mix to cool down slowly to room temperature. We then mixed 100 ng of backbone with 20 pmol of double stranded insert and performed the ligation with the Thermo Fischer Rapid DNA ligation Kit (K1422) according to manufacturer instructions. In order to obtain plasmids with an expression cassette containing two gRNAs, we ordered a 464 bp synthetic DNA fragment composed of a 430 bp sequence containing the target sequence of the first gRNA, its structural component and its terminator sequence followed by the promoter of the second gRNA and its target sequence, flanked by two LguI sites in opposite orientation (fig 1a). This 464 bp DNA fragment was digested with LguI to obtain the 430 bp insert with adequate 5’ overhangs of 3 bases and gel-purified for ligation in pGZ110. We mixed 100 ng of backbone and 20 ng of insert and performed ligation as explained previously. As an alternative to the ordering a synthetic DNA fragment, the double gRNA cassette can easily be PCR amplified using as template any plasmid already containing two gRNAs and two 55 bp oligonucleotides. The first oligo is composed of the LguI site (15 bp), the gRNA target sequence 1 (20 bp) and a homology region to the structural component of the gRNA 1 (20 bp). The second oligo is composed of the LguI site (15 bp), the gRNA target sequence 2 (20 bp) and a homology region to the promoter region of the gRNA 2 (20 bp). The resulting PCR product is then digested by LguI and cloned in the vector as previously described.

All oligonucleotides and synthetic DNA fragments were ordered from Eurofins. Refer to supplementary material for oligonucleotides and synthetic DNA sequences (supp table 1).

### Yeast transformation

Yeast cells were transformed using the standard lithium acetate method [62] with modifications. Per transformation, 10^8^ cells in Log phase were washed twice in 1 mL of double distilled water, then washed twice in 1 mL of lithium acetate mix (lithium acetate 0.1 M, TE) and cells were recovered in 50 μL of lithium acetate mix. To this mix 50 μg of denatured salmon sperm (Invitrogen), 500 ng of Cas9 plasmid, and 5 μL of double-stranded DNA donors for repair of CRISPR-induced DSBs were added. Double-stranded DNA donors for repair were prepared by mixing equimolar amounts of forward and reverse oligonucleotides at 100 pmol/µl with NEBuffer 4 (New England Biolabs), heating 5 minutes at 95°C and allowing the mix to cool down slowly to room temperature. Transformation were performed by adding 300 μL of lithium acetate/PEG solution (lithium acetate 0.1 M, PEG4000 45 %, TE), then by vortexing for 1 minute and finally incubating the cells for 25 minutes at 42°C. After transformation, cells were plated on YPD to check for viability and on synthetic medium depleted of leucine to select for transformants. Plates were incubated 4 days at 30°C. To determine the canavanine phenotypes of the transformants, 137 and 114 pink colonies from the RT and PM experiments, respectively, were resuspended in water and spot tests realized on SC and SC-arg plus canavanine (60 mg/L) agar plates. Plates were incubated 4 days at 30C before scoring the CanR/CanS phenotypes Similar tests were performed for the 7 and 30 white transformants obtained from the RT and PM experiments, respectively (fig 2B).

### Plasmid Stability

The pGZ110 plasmid is highly unstable when selection for the *LEU2* gene is removed. To cure the plasmid cell were grown overnight in YPD at 30°C. Ten individual cells were micromanipulated on YPD plates with the MSM400 micromanipulator (Singer Instruments) and grown at 30°C for 2 days. Colonies were then serially replicated on CSM-Leu and YPD. All 10 colonies lost the ability to grow on CSM-Leu.

### Estimation of CRISPR/Cas9 transformation, cutting and repair efficiencies

Efficiencies were calculated on 3 replicates of the *ADE2*/*CAN1* experiments. Transformation efficiency was defined as *p*/*T* with *p* being the average number of transformants obtained with Cas9 plasmid bearing no gRNA and without donor DNA and *T* being the average number of transformed cells. Cutting efficiency (%) was defined as 100*(*p*-*g*)/*p* with *g* being the average number of transformants obtained with Cas9 plasmid with the gRNAs and without donor DNA. Repair efficiency (%) was defined as 100\**d*/(*p*-*g*) with *d* being the average number of transformants obtained with Cas9 plasmid with the gRNAs and with donor DNAs (fig 2B).

### PFGE and colony PCR for karyotyping rearranged strains

Whole yeast chromosomes agarose plugs were prepared according to a standard method [63] and sealed in a 1% Seakem GTC agarose and 0.5x TBE gel. PFGE was conducted with the CHEF-DRII (BioRad) system with the following program: 6 V/cm for 10 hours with a switching time of 60 seconds followed by 6 V/cm for 17h with switching time of 90 seconds. The included angle was 120° for the whole duration of the run. We compared observed karyotypes with expected chromosome sizes and tested the chromosomal junctions by colony PCR with ThermoFischer DreamTaq DNA polymerase.

### Southern blot validation of *ECM34*/*SSU1* translocation

Southern blot was used to validate the translocation between *ECM34* (ch. VIII) and *SSU1* (ch. XVI). Genomic DNA was extracted from BY4741 and the engineered strain YAF082 using the Qiagen DNA buffer set (19060) and Genomic-tip 100/G (10243) according to manufacturer instructions and further purified and concentrated by isopropanol precipitation. Digestion of 10 µg of genomic DNA per strain/probe assay was carried out using FastDigest EcoRI (ThermoFischer FD0274). Electrophoresis, denaturation and neutralisation of the gel were performed according to established procedure [64]. Transfer on nylon membrane (Amersham Hybond XL) was performed using the capillarity setup [65]. The membrane was UV-crosslinked with the Stratalinker 1800 device in automatic mode. Probes targeting the genes YHL044W and *ARN1* located upstream and downstream of the cutting site in the *ECM34* promoter respectively and the genes *NOG1* and *SSU1* located upstream and downstream of the cutting site in the *SSU1* promoter respectively were amplified with ThermoFischer DreamTaq, DIG-11-dUTP deoxyribonucleotides (Roche 11 175 033 910) and gel purified. Blotting and revelation were conducted using the Roche DIG High Prime DNA Labelling and Detection Starter kit II (11 585 614 910) according to manufacturer instructions. Imaging was performed using the G:BOX Chemi XT4 (Syngene) with CSPD chemiluminescence mode. All oligonucleotides are described in supp table 1.

### Oxford Nanopore *de-novo* genome assembly

DNA from strains YAF019 and YAF064 was extracted using QIAGEN Genomic-tip 20/G columns and sheared using covaris g-TUBEs for average reads lengths of 8 kb and 15 kb respectively. DNA was repaired and dA-tailed using PreCR and FFPE kits (New England Biolabs) and cleaned with Ampure XP beads (Beckman Coulter). SQK-LSK108 adapters were ligated and libraries run on FLO-MIN107 R9.5 flowcells. Raw signals were basecalled locally using Albacore v2.0.2 with default quality filtering. Flowcell outputs are shown in supp table 2.

YAF019 was assembled using the LRSDAY v1 pipeline [66], including nanopolish v0.8.5 correction and excluding pilon polishing due to lack of illumina data. Due to only 19x coverage, the correctedErrorRate for Canu assembly was increased to 0.16. Assembly data are shown in supp table 3. YAF064 was processed using the LRSDAY v1 pipeline [66]. Linear chromosomes were assembled by SMARTdenovo v1 using 40x coverage of the longest Canu-corrected reads and combined with a Canu assembled mitochondrial genome. For canu correction, due to 200x coverage, correctedErrorRate was set at 0.75. All corrected reads were aligned against a reference-quality S288C genome, assembled with PacBio reads [67] using LAST-921. Split reads were used to highlight structural variations not apparent in the SMARTdenovo assembly. Read coverage was used to calculate an increase in the number of copies of particular regions within the rearranged genome. Evidence of rearrangements defined by overlapping reads and changes in copy number were used to manually adjust the assembly prior to nanopolish v0.8.5 and Pilon v1.22 error correction.

### Spore viability

Colonies of rearranged strains originating from the BY background and SK1 strains of the opposed mating type were mixed and spread on the same YPD plate and left overnight at 30°C. The next day, the cells were re-suspended in distilled water and single cells were picked using the Sanger MSM 400 micro-manipulator and left to grow on YPD for 2 days at 30°C until a colony appeared. Colonies originating from single cells were replicated on sporulation medium and left for 4 to 7 days at 30°C until tetrad appeared. For each cross, 10 tetrads from two diploid strains were dissected on YPD and left to grow for 3 days before counting viable spores.

### High-throughput phenotyping

Quantitative phenotyping was performed using endpoint colony growth on solid media. First, strains were grown overnight on liquid YPD medium then pinned onto a solid SC medium with a 1,536 colony per plate density format using robot assisted pinning with ROTOR™ (Singer Instrument) and incubated overnight at 30°C. Once sufficient growth is achieved, the matrix plate is replicated onto 40 media conditions (supp table 4) plus SC as a pinning control. Plates were incubated at 30°C (except for the 14°C phenotyping). After 24h, plates were imaged at a 12Mpixel resolution. Quantification of the colony size was performed in R using Gitter [68]. Raw sizes were corrected using two successive corrections: a spatial smoothing was applied to the colony size [69]. This allowed to account for variation of the plate thickness. Another correction was then applied to rescale colony size by row and column [69] which is important for colonies lying at the edges of the plate thus having easier access to nutrients compared to strains in the center. All calculations were performed using R. Once the corrected sizes were obtained, the growth ratio of each colony was computed as the colony size on the tested conditions divided by its size on SC. To detect the phenotypic effect of the engineered translocations, each growth ratio has been normalized by the growth ratio of BY4741 or BY4742, depending on the origin of the shuffled strain, on the 40 tested condition. As each strain was present six time, the value considered for its phenotype was the median of all its replicates thus smoothing pinning heterogeneity. Each experiment was repeated 2 times independently in the 40 growth conditions. Correlations between the two replicate experiments are presented in supp fig 6. For each replicate experiment and condition, the growth ratios of the 6 colonies of each tested strain were compared to the growth ratio of the reference colonies (BY4741 or BY4742) and a Wilcoxon test was used determined the significance of the phenotypic effect: ***, ** and * indicated that the two p-values from both replicates were lower than 10^-4^, 10^-3^ and 10^-2^, respectively.

## DATA ACCESS

The Oxford Nanopore sequencing data are deposited in the Sequence Read Archive under the project number (accession number pending).

## ACKNOWLEDGMENTS

We are grateful to Bruce Futcher and Gang Zhao (Stony Brook University, USA) for providing the pGZ110 plasmid. We thank our colleagues, Gianni Liti (IRCAN, France) and Bertrand Llorente (CRCM, France) for fruitful discussions and constructive suggestions as well as for providing us the wine and SK1 strains. We are also grateful to Maëlys Born-Bony for her experimental help. This work was supported by the Agence Nationale de la Recherche [ANR-16-CE12-0019]. T.F. is supported by a fellowship from the medical association la Fondation pour la Recherche Médicale. J.S. is a member of the Institut Universitaire de France.

