## Supplementary figures for "Reshuffling yeast chromosomes with CRISPR/Cas9"

```
        V 5' ATTGCGCTCTTTCCCGACGAGAGTAAATGGCGAGGATACGTTCTCTATGGAGGATGGCATAGGTGATGAAGATGAAGGAGAAGTACAGAACGCTGAAGTGAA
             |||||||||||||||||||||||||||||||||||||||||||||||||||
     VtXV 5' ATTGCGCTCTTTCCCGACGAGAGTAAATGGCGAGGATACGTTCTCTATGGATGTAGGAACATCAACATGCTCAATCTCAATCGTTAGCACATCACATTTTTC
                                                                |||||||||||||||||||||||||||||||||||||||||||||||||||
       XV 5' TGGAGAAGGGTAAATTTTTAATTTGGGATGTTTTACTTGAAGATTCTTTAGTGTAGGAACATCAACATGCTCAATCTCAATCGTTAGCACATCACATTTTTC

       XV 5' TGGAGAAGGGTAAATTTTTAATTTGGGATGTTTTACTTGAAGATTCTTTAGTGTAGGAACATCAACATGCTCAATCTCAATCGTTAGCACATCACATTTTTC
             |||||||||||||||||||||||||||||||||||||||||||||||||||
     XVtV 5' TGGAGAAGGGTAAATTTTTAATTTGGGATGTTTTACTTGAAGATTCTTTAGGGATGGCATAGGTGATGAAGATGAAGGAGAAGTACAGAACGCTGAAGTGAA
                                                                |||||||||||||||||||||||||||||||||||||||||||||||||||
        V 5' ATTGCGCTCTTTCCCGACGAGAGTAAATGGCGAGGATACGTTCTCTATGGAGGATGGCATAGGTGATGAAGATGAAGGAGAAGTACAGAACGCTGAAGTGAA

        V  5' ATTGCGCTCTTTCCCGACGAGAGTAAATGGCGAGGATACGTTCTCTATGGAGGATGGCATAGGTGATGAAGATGAAGGAGAAGTACAGAACGCTGAAGTGAA
              ||||||||||||||||||||||||||||||||||||||||||||||||||||||||||||||||||||||||||||||||||||||||||||||||||||||
restored V  5' ATTGCGCTCTTTCCCGACGAGAGTAAATGGCGAGGATACGTTCTCTATGGAGGATGGCATAGGTGATGAAGATGAAGGAGAAGTACAGAACGCTGAAGTGAA

       XV 5' TGGAGAAGGGTAAATTTTTAATTTGGGATGTTTTACTTGAAGATTCTTTAGTGTAGGAACATCAACATGCTCAATCTCAATCGTTAGCACATCACATTTTTC
              ||||||||||||||||||||||||||||||||||||||||||||||||||||||||||||||||||||||||||||||||||||||||||||||||||||||
restored XV 5' TGGAGAAGGGTAAATTTTTAATTTGGGATGTTTTACTTGAAGATTCTTTAGTGTAGGAACATCAACATGCTCAATCTCAATCGTTAGCACATCACATTTTTC
```

**Supplementary figure 1** Sanger sequencing of the translocated junctions in YAF190, YAF192 and de-translocated junctions in strains YAF194, YAF199. The sequences corresponding to donor oligonucleotides are shown in bold. The two gRNA target sequences are highlighted in light blue and orange. PAM sequences are highlighted in dark blue and orange.

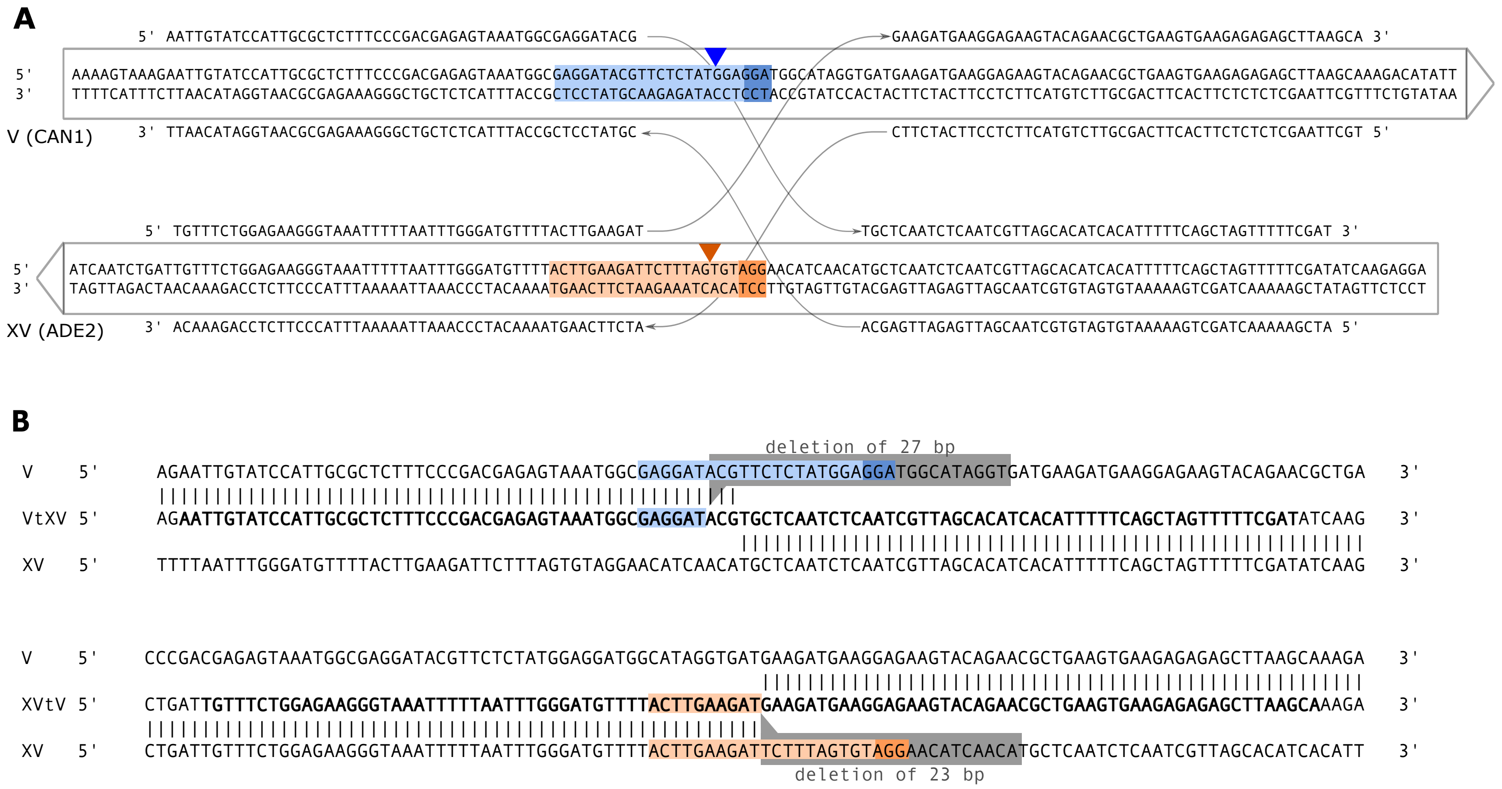


**Supplementary figure 2 (A)** Targeted sequences and donors used to engineer the translocation with a small deletion. The two gRNA target sequences are highlighted in light blue and orange. PAM sequences are highlighted in dark blue and orange. Triangles indicate DSBs sites. Arrows framing the sequences indicate the orientation of coding phases. Donor nucleotides are represented above and below the frames by sequences linked by thin arrows to indicate their homology with the two different chromosomes. **(B)** Alignments of the *de-novo* assembled chimerical junctions on reference chromosomes V and XV. Donor sequences used to direct the translocation are in bold. Deleted sequences in chromosomes V and XV are highlighted in grey. The translocation occurred at the targeted position with a base-pair resolution.

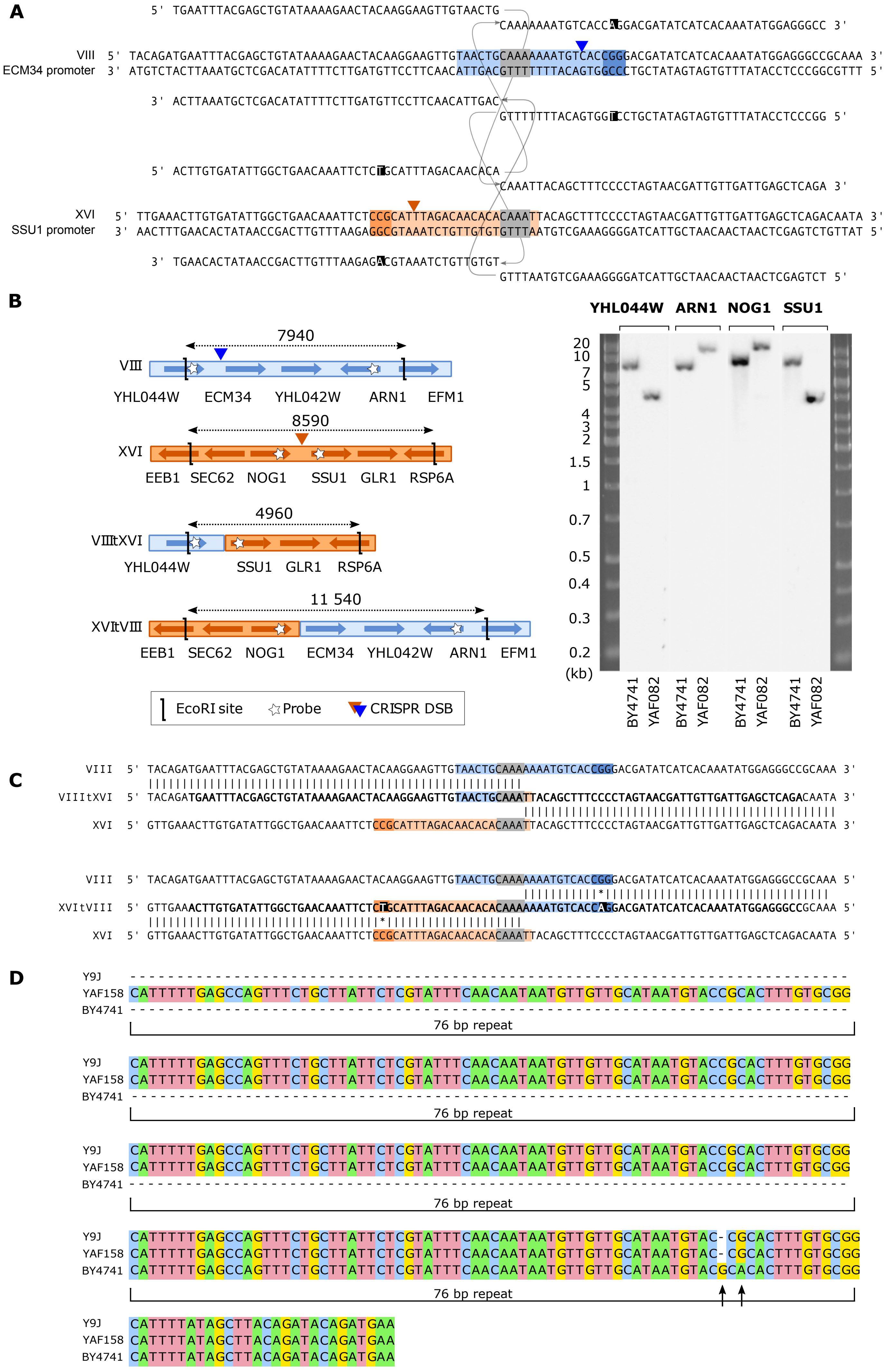

A
5' TGAATTTACGAGCTGTATAAAAGAACTACAAGGAAGTTGTAACTG
CAAAAAAATGTCACCAGGACGATATCATCACAAATATGGAGGGCC 3'
VIII
ECM34 promoter
5' TACAGATGAATTTACGAGCTGTATAAAAGAACTACAAGGAAGTTGTAACTGCAAAAAAATGTCACCGGGACGATATCATCACAAATATGGAGGGCCGCAAA 3'
3' ATGTCTACTTAAATGCTCGACATATTTTCTTGATGTTCCTTCAACATTGACGTTTTTTTACAGTGGCCCTGCTATAGTAGTGTTTATACCTCCCGGCGTTT 5'
3' ACTTAAATGCTCGACATATTTTCTTGATGTTCCTTCAACATTGAC
GTTTTTTTACAGTGGTCCTGCTATAGTAGTGTTTATACCTCCCGG 5'
5' ACTTGTGATATTGGCTGAACAAATTCTCTGCATTTAGACAACACA
CAAATTACAGCTTTCCCCTAGTAACGATTGTTGATTGAGCTCAGA 3'
XVI
SSU1 promoter
5' TTGAAACTTGTGATATTGGCTGAACAAATTCTCCGCATTTAGACAACACACAAATTACAGCTTTCCCCTAGTAACGATTGTTGATTGAGCTCAGACAATA 3'
3' AACTTTGAACACTATAACCGACTTGTTTAAGAGGCGTAAATCTGTTGTGTGTTTAATGTCGAAAGGGGATCATTGCTAACAACTAACTCGAGTCTGTTAT 5'
3' TGAACACTATAACCGACTTGTTTAAGAGACGTAAATCTGTTGTGT
GTTTAATGTCGAAAGGGGATCATTGCTAACAACTAACTCGAGTCT 5'
B
7940
VIII
YHL044W ECM34 YHL042W ARN1 EFM1
8590
XVI
EEB1 SEC62 NOG1 SSU1 GLR1 RSP6A
4960
VIIItXVI
YHL044W SSU1 GLR1 RSP6A
11 540
XVItVIII
EEB1 SEC62 NOG1 ECM34 YHL042W ARN1 EFM1
EcoRI site
Probe
CRISPR DSB
YHL044W ARN1 NOG1 SSU1
20
10
7
5
4
3
2
1.5
1
0.7
0.5
0.4
0.3
0.2
(kb)
BY4741 YAF082 BY4741 YAF082 BY4741 YAF082 BY4741 YAF082
C
VIII 5' TACAGATGAATTTACGAGCTGTATAAAAGAACTACAAGGAAGTTGTAACTGCAAAAAAATGTCACCGGGACGATATCATCACAAATATGGAGGGCCGCAAA 3'
VIIItXVI 5' TACAGATGAATTTACGAGCTGTATAAAAGAACTACAAGGAAGTTGTAACTGCAAATTACAGCTTTCCCCTAGTAACGATTGTTGATTGAGCTCAGACAATA 3'
XVI 5' GTTGAAACTTGTGATATTGGCTGAACAAATTCTCCGCATTTAGACAACACACAAATTACAGCTTTCCCCTAGTAACGATTGTTGATTGAGCTCAGACAATA 3'
VIII 5' TACAGATGAATTTACGAGCTGTATAAAAGAACTACAAGGAAGTTGTAACTGCAAAAAAATGTCACCGGGACGATATCATCACAAATATGGAGGGCCGCAAA 5'
XVItVIII 5' GTTGAAACTTGTGATATTGGCTGAACAAATTCTCTGCATTTAGACAACACACAAAAAAATGTCACCAGGACGATATCATCACAAATATGGAGGGCCGCAAA 5'
XVI 5' GTTGAAACTTGTGATATTGGCTGAACAAATTCTCCGCATTTAGACAACACACAAATTACAGCTTTCCCCTAGTAACGATTGTTGATTGAGCTCAGACAATA 5'
D
Y9J
YAF158 CATTTTTGAGCCAGTTTCTGCTTATTCTCGTATTTCAACAATAATGTTGTTGCATAATGTACCGCACTTTGTGCGG
BY4741
76 bp repeat
Y9J CATTTTTGAGCCAGTTTCTGCTTATTCTCGTATTTCAACAATAATGTTGTTGCATAATGTACCGCACTTTGTGCGG
YAF158 CATTTTTGAGCCAGTTTCTGCTTATTCTCGTATTTCAACAATAATGTTGTTGCATAATGTACCGCACTTTGTGCGG
BY4741
76 bp repeat
Y9J CATTTTTGAGCCAGTTTCTGCTTATTCTCGTATTTCAACAATAATGTTGTTGCATAATGTACCGCACTTTGTGCGG
YAF158 CATTTTTGAGCCAGTTTCTGCTTATTCTCGTATTTCAACAATAATGTTGTTGCATAATGTACCGCACTTTGTGCGG
BY4741
76 bp repeat
Y9J CATTTTTGAGCCAGTTTCTGCTTATTCTCGTATTTCAACAATAATGTTGTTGCATAATGTAC-CGCACTTTGTGCGG
YAF158 CATTTTTGAGCCAGTTTCTGCTTATTCTCGTATTTCAACAATAATGTTGTTGCATAATGTAC-CGCACTTTGTGCGG
BY4741 CATTTTTGAGCCAGTTTCTGCTTATTCTCGTATTTCAACAATAATGTTGTTGCATAATGTACGCACACTTTGTGCGG
76 bp repeat
Y9J CATTTTATAGCTTACAGATACAGATGAA
YAF158 CATTTTATAGCTTACAGATACAGATGAA
BY4741 CATTTTATAGCTTACAGATACAGATGAA

**Supplementary figure 3: Engineering the SSU1-ECM34 translocation. A.** gRNAs and donors used to perform the translocation between the promoters of SSU1 and ECM34. The micro-homology regions between the two chromosomes are highlighted in grey. The two gRNA target sequences are highlighted in light blue and orange. PAM sequences are highlighted in dark blue and orange. Triangles indicate DSBs sites. Mismatches brought by the donors to inactivate the PAMs are highlighted in black. **B.** Validation of the SSU1-ECM34 translocation by southern blot. Left: schematic view of the chromosomal regions surrounding the translocation breakpoints. The double arrows indicate the length of the EcoRI restriction fragments. Right: Southern blot on the translocated strain YAF082 and parental strain BY4741. The probes are indicated on the top of the lanes. **C.** Sanger sequencing of YAF82 chimerical junctions. The two mismatches brought by the donors to prevent CRISPR/Cas9 activity after repair (indicated by stars) are present in chromosome XVItVIII. **D.** Multiple alignment of the promoter regions in front of the SSU1 gene in Y9J and YAF158 and of ECM34 in BY4741. Note that there are only 3 copies of the repeats present in the published assembly of the Y9J strain while it actually contains 4 repeats because the PCR product inserted in the YAF158 strain was amplified using the Y9J genome as matrix. There is only 1 copy of the 76 repeat region in the BY4741 genome. The 2 black arrows point to one indel and 1 SNP between the sequence of the BY4741 and the 2 other strains.

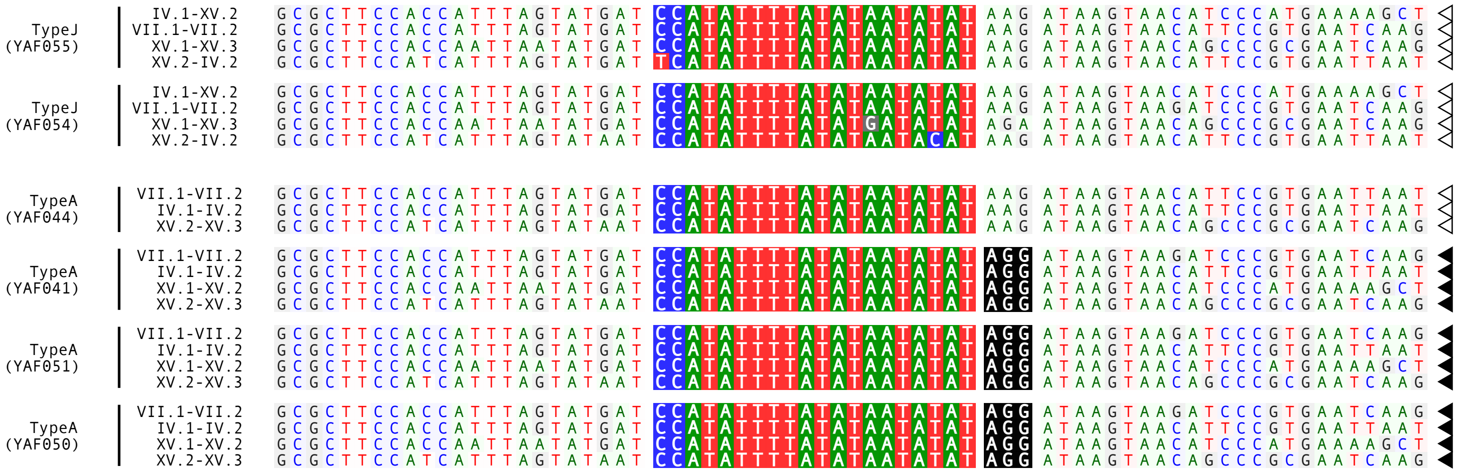

**Supplementary figure 4** Sanger sequencing of the junctions of two type J strains (YAF055 and YAF054) and four type A strains (YAF044, YAF041, YAF051, YAF050). Junctions where the PAM was mutated are indicated by white triangles. Sequences where the PAM is intact are indicated by black triangles and PAM is highlighted in black.

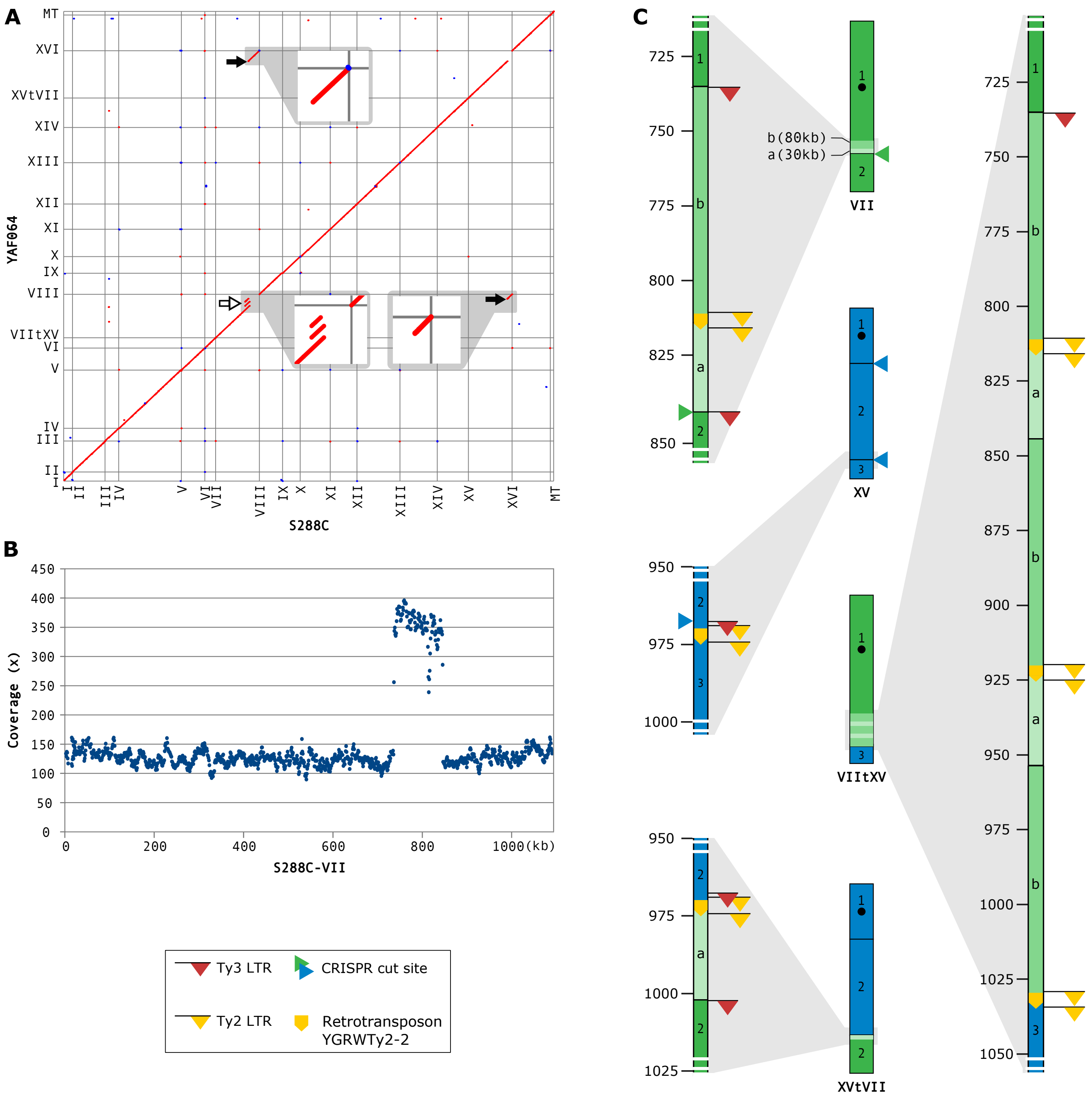


**Supplementary figure 5: Genome with complex rearrangement. A.** Homology matrix between the genomes of the strain showing an increase in global DNA content in PFGE (YAF064) and S288c. Translocated fragments are indicated by black arrows. The tandem triplication at the junction of chromosome VIItXV is indicated by the white arrow. **B.** Coverage of the YAF064 reads remapped on the reference chromosome VII. Each dot represents a window of 1kb. **C.** Architecture of chromosomes VII (in green) and XV (in blue) of the reference strain and chimerical chromosomes VIItXV and XVtVII of the rearranged strain YAF064. Light grey triangles represent zoom-ins on chromosomal junctions. Ty3-LTRs and Ty2 LTRs elements are represented by red and yellow flags respectively. Full-length Ty2 elements are represented by yellow boxes. The Ty3 LTR copies targeted by CRISPR/Cas9 are indicated by green and blue triangles next to chromosomes VII and XV, respectively. The displaced 30kb segment is referred to as region a and the other 80 kb segment as region b. Regions a and b, triplicated in the shuffled strain, are represented in lighter green shades. In summary, one copy of region a lies at the chimerical junction of chromosome XVtVII, whereas the remaining two and three copies of region a and b, respectively, are found in tandem at the chimerical junction of chromosome VIItXV.

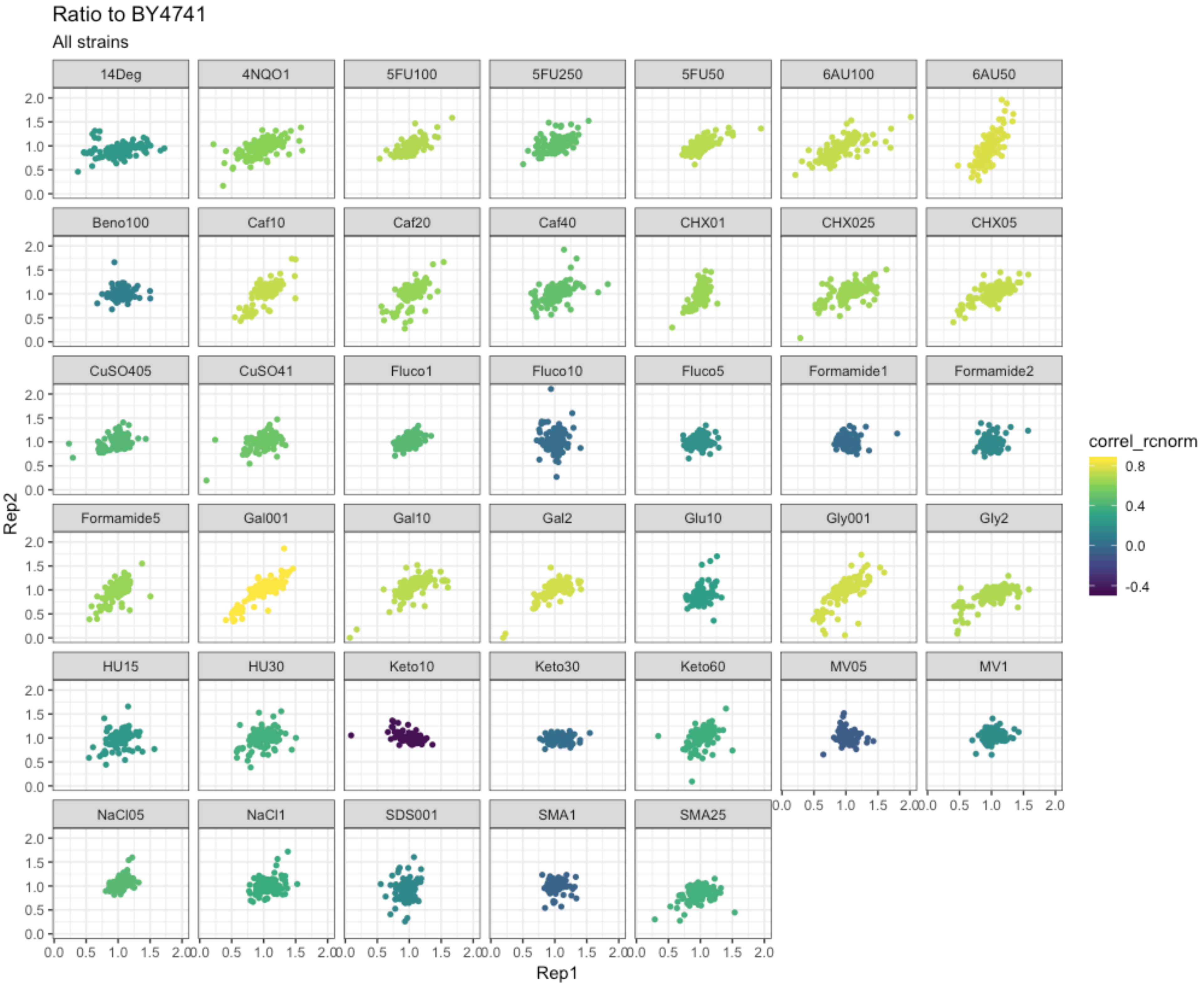

Ratio to BY4741
All strains
14Deg
4NQO1
5FU100
5FU250
5FU50
6AU100
6AU50
Beno100
Caf10
Caf20
Caf40
CHX01
CHX025
CHX05
CuSO405
CuSO41
Fluco1
Fluco10
Fluco5
Formamide1
Formamide2
Formamide5
Gal001
Gal10
Gal2
Glu10
Gly001
Gly2
HU15
HU30
Keto10
Keto30
Keto60
MV05
MV1
NaCl05
NaCl1
SDS001
SMA1
SMA25
Rep2
Rep1
correl_rcnorm
0.8
0.4
0.0
-0.4

**Supplementary figure 6: high throughput phenotyping, correlations between the two replicate experiments.** Each plot represents one condition and each dot represents the growth ratio of each strain (i.e. the colony size on the tested conditions divided by its size on SC) divided by the growth ration of BY4741.
